# Postnatal Tendon Growth and Remodeling Requires Platelet-Derived Growth Factor Receptor Signaling

**DOI:** 10.1101/208991

**Authors:** Kristoffer B Sugg, James F Markworth, Nathaniel P Disser, Andrew M Rizzi, Jeffrey R Talarek, Dylan C Sarver, Susan V Brooks, Christopher L Mendias

**Author notes:** To whom correspondence should be addressed: Christopher L Mendias, Hospital for Special Surgery, 535 E 70th St, New York, NY 10021.

## Abstract

Platelet-derived growth factor receptor (PDGFR) signaling plays an important role in the fundamental biological activities of many cells that compose musculoskeletal tissues. However, little is known about the role of PDGFR signaling during tendon growth and remodeling in adult animals. Using the hindlimb synergist ablation model of tendon growth, our objectives were to determine the role of PDGFR signaling in the adaptation of tendons subjected to a mechanical growth stimulus, as well as to investigate the biological mechanisms behind this response. We demonstrate that both PDGFRs, PDGFRα and PDGFRβ, are expressed in tendon fibroblasts, and that the inhibition of PDGFR signaling suppresses the normal growth of tendon tissue in response to mechanical growth cues due to defects in fibroblast proliferation and migration. We also identify that membrane type-1 matrix metalloproteinase (MT1-MMP) as an essential proteinase for the migration of tendon fibroblasts through their extracellular matrix. Furthermore, we report that MT1-MMP translation is regulated by PI3K/Akt signaling, while ERK1/2 controls post-translational trafficking of MT1-MMP to the plasma membrane of tendon fibroblasts. Taken together, these findings demonstrate that PDGFR signaling is necessary for postnatal tendon growth and remodeling, and that MT1-MMP is a critical mediator of tendon fibroblast migration and a potential target for the treatment of tendon injuries and diseases.

## INTRODUCTION

Tendon is an integral component of the musculoskeletal system. Anatomically situated between skeletal muscle and bone, tendon transmits and stores force which allows for efficient locomotion. The function of tendon is determined by the biochemical composition and macromolecular structural organization of its extracellular matrix (ECM), which consists of a dense network of cross-linked type I collagen and smaller amounts of type III collagen, elastin and various proteoglycans (27). This collagen-rich tissue provides structural support to the tendon, selectively binds and releases growth factors that regulate multiple cellular functions, and organizes the compartments that contain various cell populations (9). Tendon fibroblasts are the predominant cell type in tendon, and are responsible for the production, maintenance, modification and repair of matrix proteins (27). Despite the importance of tendon to the overall function of the musculoskeletal system, relatively little is known about the cellular and molecular mechanisms that regulate tendon growth and remodeling in adult animals.

Tendon adapts to increased mechanical signals by undergoing hypertrophy, as evidenced by increases in tendon cross-sectional area (CSA), cell density, peak stress, peak strain and type I collagen content in response to loading (6, 14, 34). While the specific growth factors and signaling pathways required for this process are largely unknown, one family of growth factors that are upregulated following tendon injury and are active at multiple stages of the healing process are the platelet-derived growth factors (PDGFs) (35). PDGFs typically function as secreted hetero- or homodimers of disulfide-linked polypeptide chains (PDGF-AA, PDGF-BB, PDGF-CC, PDGF-DD and PDGF-AB) (1), with PDGF-BB among the most frequent PDGF isoforms used to enhance regeneration in various animal models of tendon injury (29, 57). PDGFs bind to and activate a class of structurally related receptor tyrosine kinase transmembrane proteins known as PDGF receptors α (PDGFRα) and β (PDGFRβ) (1). Upon binding to their ligands, PDGFRs dimerize and undergo autophosphorylation of conserved cytoplasmic tyrosine residues that initiate multiple signal transduction cascades, including the phosphoinositide 3-kinase (PI3K)/protein kinase B (Akt) and extracellular signal-related kinases 1 and 2 (ERK1/2) pathways (1, 52). PDGFs are potent mitogens and their signaling events control key biological functions in many cell types of mesenchymal origin, including proliferation, differentiation, migration and ECM synthesis and remodeling (50). While PDGFs are expressed in tendon tissue, and active at multiple stages of the healing process in injured tendons (35), the identity and tissue localization of PDGFRα^+^ and PDGFRβ^+^ cells in tendon have not been clearly defined, and the overall importance of PDGFR signaling to the growth response of mechanically loaded tendons is presently unknown.

During periods of tendon growth and remodeling, fibroblasts express matrix metalloproteinases (MMPs), a family of zinc-dependent endopeptidases that collectively degrade multiple ECM components, including type I collagen (9, 48). Membrane-type MMPs (MT-MMPs) are a sub-class of MMPs that allow cells to migrate through their respective ECMs, by constraining type I collagen degradation to the leading edge of the cell membrane (20, 45).
Within the MT-MMP subgroup, membrane type-1 MMP (MT1-MMP) is required for proper tendon development, as the deletion of MT1-MMP in embryonic tendon fibroblasts results in the failure of normal tendon formation (55). MT1-MMP expression has also been shown to correlate with pivotal events in the growth response of mechanically-loaded tendons of adult animals (48). In other cell types, MT1-MMP expression is regulated by receptor tyrosine kinase signaling (19, 52), but the signal transduction pathways that regulate MT1-MMP expression in tendon fibroblasts are not known.

Given that PDGFs are expressed in tendon, and that MT1-MMP is important for tendon development, we sought to determine the role of PDGFR signaling in postnatal tendon growth and remodeling. We hypothesized that inhibition of PDGFR signaling would suppress the normal growth of tendon tissue during mechanical overload due to defects in cell proliferation and migration. To test this hypothesis, we induced tendon growth in adult mice via mechanical overload using the hindlimb synergist ablation model which has been previously used to study muscle and tendon growth, and blocked PDGFRα and PDGFRβ signaling through the use of a highly specific pharmacological inhibitor CP-673,451. We also conducted a series of *in vitro* experiments to determine the molecular mechanisms that underlie the PDGFR-dependent growth of tendons in adult animals.

## METHODS

*Mice.* All animal studies were approved by the University of Michigan Institutional Animal Care & Use Committee. Wild-type (WT) C57BL/6 mice and transgenic *PDGFRα*^*eGFP/*+^ mice were obtained from the Jackson Laboratory (Bar Harbor, ME, USA). *PDGFRα*^*eGFP/*+^ mice express the nuclear-localized H2B-eGFP reporter gene from the endogenous PDGFRa locus (17). Four-month-old male mice were used in all experiments.

*Synergist Ablation.* Mice were randomized to 3- and 10-day groups. Bilateral synergist ablation procedures were performed under isoflurane anesthesia as described previously (14, 50). An overview is presented in Figure 1A. The Achilles tendon was surgically excised to prevent the gastrocnemius and soleus muscles from plantarflexing the talocrural joint, resulting in compensatory hypertrophy of the synergist plantaris muscle-tendon unit. Buprenorphine was administered for post-operative analgesia, and *ad libitum* weight-bearing and cage activity were allowed in the postoperative period. Mice were closely monitored during the postoperative period for any adverse reactions. At tissue harvest, the left plantaris tendons were collected for gene expression analysis, while the right plantaris tendons were used for histological examination. After the tendons were removed, mice were euthanized by cervical dislocation and induction of bilateral pneumothorax. Plantaris tendons from additional non-overloaded mice were obtained as described above for gene expression analysis.

**Figure 1.**
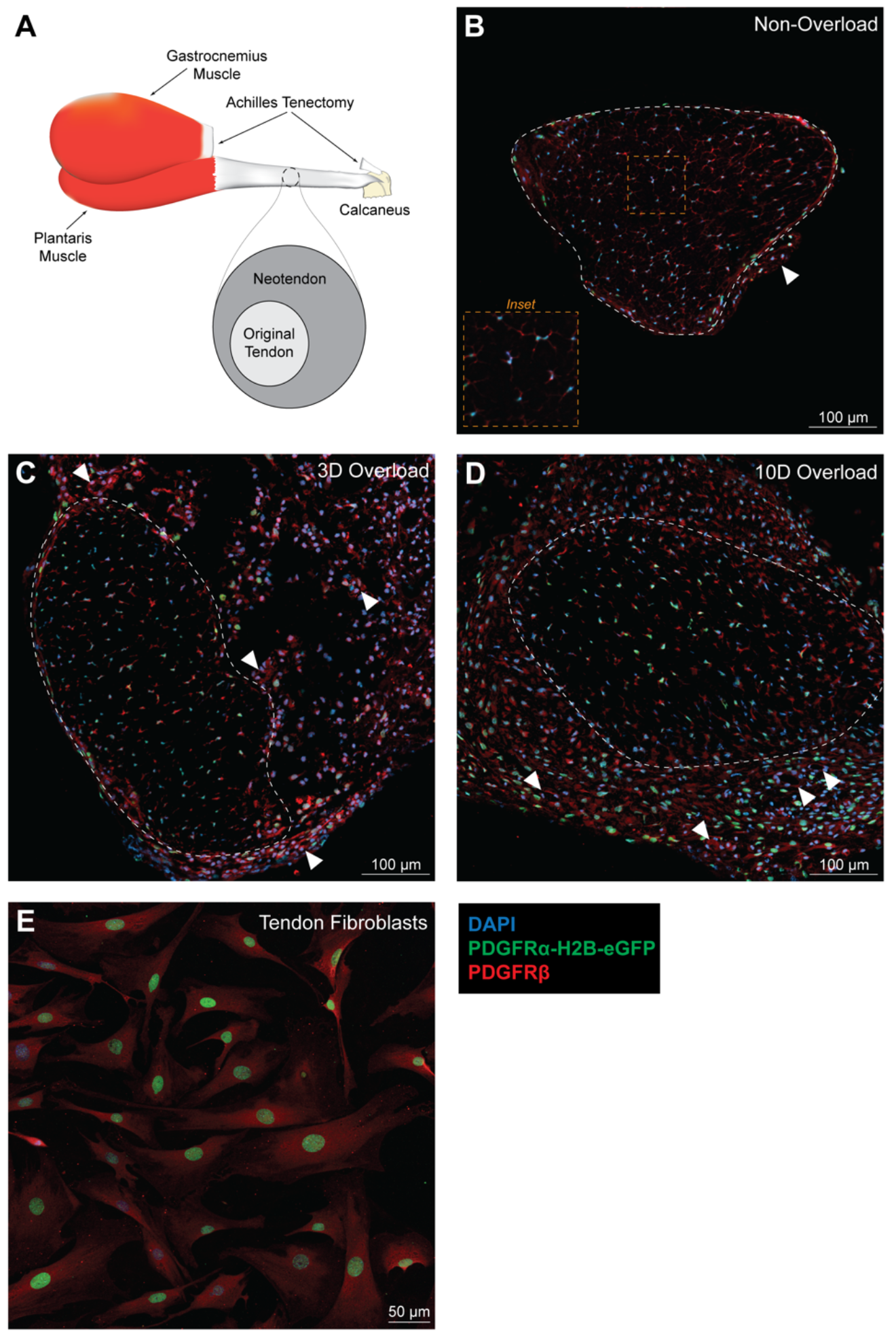
Tendon fibroblasts express both PDGFRα and PDGFRβ. (A) Schematic of the synergist ablation procedure. All histological sections were taken from the midportion of the plantaris tendon. Representative images of (B) non-overloaded, (C) 3-day overloaded and (D) 10-day overloaded plantaris tendons. Dashed white line indicates the boundary between the original tendon and neotendon, and a dashed yellow line high magnification inset is provided for Figure 1B. White arrowheads indicate blood vessels composed of PDGFRα^−^/PDGFRβ^+^ cells. Scale bars are 100 μm. (E) Tendon fibroblasts were isolated from *PDGFRα*^*eGFP/*+^ mice and immunostained for PDGFRβ. Scale bar is 50 μm. DAPI, blue; PDGFRα-H2B-eGFP, green; PDGFRβ, red.

*Inhibition of PDGFR Signaling.* Within each group of mice, half of the mice were treated with vehicle and half with CP-673,451 (Biorbyt, Cambridge, UK), a specific inhibitor of PDGFRα and PDGFRβ (10, 43). CP-673,451 was dissolved first in 2 parts dimethyl sulfoxide and then in 8 parts phosphate-buffered saline. CP-673,451 was administered by intraperitoneal injection at 15 mg/kg twice daily, with a total daily dose of 30 mg/kg, starting the day prior to synergist ablation and continued each day until tissue harvest. This dose has previously been shown to be effective at inhibiting PDGFR phosphorylation in mice *in vivo* (43, 50).

*Histology.* Histological examination of the tendon tissue was performed as described previously (48, 50). Plantaris tendons were placed in 30% sucrose solution for one hour, snap frozen in Tissue-Tek OCT Compound (Sakura Finetek, Torrance, CA, USA) and stored at −80°C until use. Tendons were sectioned at a thickness of 10 μm in a cryostat and stained with hematoxylin and eosin (H&E) to determine tendon CSA and cell density. For immunohistochemistry, tendon sections were fixed in 4% paraformaldehyde, permeabilized with 0.2% Triton X-100, and blocked with 5% goat serum. To identify cell types in non-overloaded and overloaded tendons, slides from *PDGFRPα*^*EGFP/*+^ mice were incubated with rabbit anti-PDGFRβ (1:100; sc-339, Santa Cruz Biotechnology, Santa Cruz, CA, USA) primary antibodies. Proliferating cells were identified in overloaded tendons treated with vehicle or PDGFR inhibitor, by incubating slides from WT mice subjected to synergist ablation with rabbit anti-Ki67 (1:100; ab16667, Abcam, Cambridge, MA, USA) primary antibodies. Primary rabbit antibodies against p-PDGFRα^Y849^/PDGFRβ^Y857^ (1:100; #3170, Cell Signaling Technology, Danvers, MA, USA) were also used on slides from C57BL/6 mice to determine differences in the level of PDGFR phosphorylation in response to vehicle or PDGFR inhibitor treatment. The ECM was identified with wheat germ agglutinin (WGA) lectin conjugated to Alexa Fluor 488 (AF488) (1:200; W11261, Thermo Fisher Scientific, Carlsbad, CA, USA). Secondary antibodies conjugated to AF555 (1:300; A-21429, Thermo Fisher Scientific) were used to detect primary antibodies. Nuclei were stained with DAPI (1:500; D9542, Sigma Aldrich, St. Louis, MO, USA). High-resolution digital images were captured with an Olympus BX-51 microscope and camera (Olympus, Center Valley, PA, USA) for the H&E and Ki67 slides, while a Nikon A1 confocal laser microscrope (Nikon Instruments, Tokyo, JP) was used for the PDGFR slides. Quantification was performed in a blinded fashion using ImageJ software (NIH, Bethesda, MD, USA).

*Cell Culture.* Fibroblasts were isolated from the tail tendons of mice as described previously (22). Tail tendons are useful for obtaining a large number of early passage cells, and they arise from the same population of somitic progenitor cells as limb tendons during development (36). Briefly, mice were anesthetized as described above, the tail was removed, and animals were euthanized by cervical dislocation. Fascicles were isolated from tail tendons, and then were finely minced with scissors and placed in low-glucose Dulbecco’s Modified Eagle Medium (DMEM; Gibco, Carlsbad, CA, USA) containing 0.2% type II collagenase (Gibco) for 1 hour at 37°C with constant agitation. An equal volume of growth medium (GM) that consists of low-glucose DMEM with 10% fetal bovine serum (FBS; Gibco) and 1% antibiotic-antimycotic (Gibco) was added to the digested tissue to inactivate the collagenase. Cells were pelleted by centrifugation at 300*g* for 10 minutes, resuspended in GM and plated on 100-mm type I collagen-coated dishes (Corning, Corning, NY, USA). All cells were maintained in humidified incubators at 37°C and 5% CO_2_. Passage 2-3 cells were used in all experiments. To block PDGFR signaling, cells were incubated with 1 μM CP-673,451, as described above.

*Migration Assay.* Type I collagen was acid-extracted from cadaver rat tail tendons as described (20). Briefly, tendons were excised and placed in 0.2% acetic acid for 5 days at 4°C. The collagen solution was then centrifuged at 24,000*g* for 30 minutes, and the supernatant was collected, lyophilized and dissolved again in 0.2% acetic acid to a final concentration of 2.7 mg/ml. Collagen gels were prepared in the upper chambers of Transwell dishes (12-mm diameter, 3-μm pore size; Corning) by combining the collagen solution with 10× Minimum Essential Medium (MEM, Thermo Fisher Scientific) and 0.34 N NaOH in an 8:1:1 ratio. After 45 minutes at 37°C, the gelling process was complete and 1×10^5^ tendon fibroblasts were seeded on top of the collagen gels. Growth factors, either PDGF-BB (20 ng/ml; R&D Systems, Minneapolis, MN, USA) or TGF-β1 (10 ng/ml; PeproTech, Rocky Hill, NJ, USA), were added to the lower chambers. Where indicated, media was supplemented with the synthetic broad spectrum MMP inhibitor BB-94 (5 μM; Tocris Bioscience, Bristol, UK), the mitogen-activated protein kinase kinases 1 and 2 (MEK1/2) inhibitor PD98059 (50 μM; InvivoGen, San Diego, CA, USA) or the PI3K inhibitor wortmannin (10 μM; InvivoGen). All media, including growth factors and inhibitors, were replaced every 2 days. Migratory activity was monitored by phase-contrast microscopy, and the cells were allowed to migrate for 6 days. At the completion of the experiment, gels were fixed with 10% neutral buffered formalin, embedded in paraffin, sectioned at a thickness of 10μm, and stained with hematoxylin and eosin. The number of migrating cells and the maximum distance migrated were quantified in 5 randomly selected fields of a single experiment from 3 or more independent experiments performed.

*Proliferation Assay.* Uptake of bromodeoxyuridine (BrdU) by proliferating tendon fibroblasts was measured as described previously (33). Tendon fibroblasts were incubated overnight with or without 20 ng/ml of PDGF-BB in media containing 0.5% FBS. After overnight incubation, fresh media was added along with 20 μM of BrdU (Sigma Aldrich) for 1 hour. Cells were then rinsed with phosphate-buffered saline, fixed in ice-cold methanol and permeabilized with 0.5% Triton X-100. The BrdU epitope was exposed by denaturing DNA with 2 M HCl. BrdU was visualized with an anti-BrdU antibody (1:50; G3G4, Developmental Studies Hybridoma Bank, Iowa City, Iowa, USA) and a secondary antibody conjugated to AF555 (1:200; A-21127, Thermo Fisher Scientific). Nuclei were counterstained with DAPI (1:500; D9542, Sigma Aldrich) to determine total cell number. The number of proliferating cells were quantified in five randomly selected fields of a single experiment from 6 independent experiments performed.

*Quantitative RT-PCR.* Gene expression analysis was performed as described previously (22, 51). Plantaris tendons were homogenized in QIAzol (Qiagen, Valencia, CA, USA) and RNA was purified using a miRNeasy Micro Kit (Qiagen) supplemented with DNase I (Qiagen). RNA was reverse transcribed into cDNA with iScript Reverse Transcription Supermix (Bio-Rad, Hercules, CA, USA). Amplification of cDNA was performed in a CFX96 real-time thermal cycler (Bio-Rad) using iTaq Universal SYBR Green Supermix (Bio-Rad). Target gene expression was normalized to the stable housekeeping gene peptidylprolyl isomerase D (PPID), and further normalized to tendons that were not subjected to synergist ablation using the 2^−ΔΔCt^ method. PPID was selected as a housekeeping gene from microarray data and validated with qPCR. For cell culture experiments, relative transcript copy number was calculated using the linear regression of efficiency method (46). Primer sequences are provided in Table S1.

*Microarray*. Microarray measurements were performed by the University of Michigan DNA Sequencing Core as described previously (22, 50). Equal amounts of RNA isolated from four individual tendons were pooled into a single sample for microarray analysis, and two pooled samples from each group were analyzed. RNA was pooled because gene expression from a pooled sample is similar to the average of the individual samples composing the pooled sample (5, 25). Biotinylated cDNA was prepared using the GeneChip WT PLUS Reagent Kit (Affymetrix, Santa Clara, CA, USA) and hybridized to Mouse Gene 2.1 ST Array Strips (Affymetrix). Raw microarray data were loaded into ArrayStar version 12.1 (DNASTAR, Madison, WI, USA) to calculate fold changes in gene expression. The microarray dataset has been deposited to the NIH Gene Expression Omnibus database (accession number GSE95794).

*siRNA Transfection.* Predesigned fluorescent-labeled siRNAs directed against mouse MT1-MMP (NM_008608; SI02733822, Qiagen) were transfected into tendon fibroblasts using Lipofectamine RNAiMAX (Thermo Fisher Scientific) at a final concentration of 10 nM. AllStars Negative Control siRNA (Qiagen) was used as a negative control.

*Immunoblots.* Immunoblots were performed as described (22, 50). Whole tendons and cell pellets were homogenized in ice cold RIPA buffer (Sigma Aldrich) supplemented with 1% protease and phosphatase inhibitor cocktail (Thermo Fisher Scientific). Protein homogenates were diluted in Laemmli’s sample buffer, boiled for two minutes and 20 μg of protein was separated on either 6% or 12% SDS-PAGE gels depending on the protein of interest. Proteins were transferred to 0.45-μm nitrocellulose membranes (Bio-Rad) using the Trans-Blot SD semi-dry transfer apparatus (Bio-Rad), blocked with 5% non-fat powdered milk in TBST solution and incubated with primary rabbit antibodies (1:1000; Cell Signaling Technology) against p-PDGFRα^Y849^/PDGFRβ^Y857^ (#3170), PDGFRα (#3174), PDGFRβ (#3169), p-ERK1/2^T202/Y204^ (#4370), ERK1/2 (#4695), p-Akt^T308^ (#13038), Akt (#4691), p-p70S6K^T389^ (#9234) and p70S6K (#2708), primary rabbit antibodies (1:1000; Abcam) against MT1-MMP (ab51074) or primary rabbit antibodies (1:1000; Santa Cruz Biotechnology) against procollagen type I (Pro-Col1a1; sc-30136). β-tubulin (ab6046, Abcam) or Coomassie staining were used to determine equal protein loading. Following primary antibody incubation, membranes were rinsed and incubated with HRP-conjugated goat anti-rabbit secondary antibodies (1:10,000; ab97051, Abcam). Proteins were detected using enhanced chemiluminescent reagents (Bio-Rad) and visualized using a digital chemiluminescent documentation system (Bio-Rad), and expressed as arbitrary densitometry units (AU).

*Cell Surface Biotinylation.* Cell surface proteins from tendon fibroblasts were biotinylated and purified using the Cell Surface Protein Isolation Kit (Thermo Fisher Scientific). Briefly, cells in monolayer were washed with ice-cold phosphate-buffered saline and incubated with 0.25 mg/ml of Sulfo-NHS-SS-Biotin for 30 minutes at 4 °C with constant agitation. Adherent cells were then lysed and the biotinylated proteins were affinity-purified using streptavidin agarose beads. Coomassie staining was used to determine equal protein loading for immunoblots. Biotinylated proteins were analyzed by immunoblotting as described above with anti-MT1-MMP (ab51074, Abcam) antibodies, while GAPDH (MA5-15738, Thermo Fisher Scientific) served as the control for cytosolic proteins.

*Statistics.* Results are presented as mean±SD. Prism version 7.0 (GraphPad Software, La Jolla, CA, USA) was used to conduct statistical analyses. A two-way ANOVA (α=0.05) followed by Tukey’s post hoc sorting evaluated the interaction between time after synergist ablation and PDGFR inhibitor treatment. For cell culture experiments, differences between groups were tested with an unpaired Student’s t-test or a one-way ANOVA followed by Tukey’s post hoc sorting where indicated with α=0.05.

## RESULTS AND DISCUSSION

*PDGFRα and PDGFRβ are expressed in tendon fibroblasts.* We first identified which cells express the PDGFRs in tendon tissue (Figure 1A-D). Tendon fibroblasts, which are the major cell population in tendon, express either PDGFRα or PDGFRβ, and in some cases both receptors (Figure 1B). We then performed mechanical overload of the plantaris muscle-tendon unit by removal of the tendon which transmits the load from the synergist gastrocnemius and soleus muscles. This techniques has been used to study muscle hypertrophy in numerous studies (11, 16, 32, 50), and we and others have also previously used this to study tendon hypertrophy (14, 39, 48). In this model, the neotendon tissue that forms around the existing tendon is populated by proliferating tendon fibroblasts, that mature and generate a mature ECM that resembles the original tendon matrix (14, 39, 48). After mechanical overload, we also observed a substantial number of PDGFRα^+^/PDGFRβ^+^ cells in the neotendon tissue (Figure 1C-D). Very few cells were found that express PDGFRα alone (PDGFRα^+^/PDGFRβ^−^), and a small population of cells near blood vessels expressed PDGFRβ alone (PDGFRα^−^/PDGFRβ^+^). *In vitro*, tendon fibroblasts express both PDGFR subtypes (PDGFRα^+^/PDGFRβ^+^) (Figure 1E). That most of the fibroblasts in tendon express PDGFRα and PDGFRβ, with some cells demonstrating expression of both receptors, differs from what is observed in the extracellular matrix of skeletal muscle where populations of PDGFR-expressing cells can be separated into two distinct groups. PDGFRα is expressed by fibroadipogenic precursor cells or fibroblasts in the skeletal muscle interstitium, and PDGFRβ is expressed by perivascular cells or pericytes that often behave as pluripotent stem cells, with almost no cells demonstrating expression of both PDGFR isoforms (24, 50, 58). While tendon has PDGFRα^−^/PDGFRβ^+^ expressing cells around blood vessels that are likely pericytes, even though the ECM of skeletal muscle and tendon are intimately linked, the finding that most tendon fibroblasts express PDGFRα and PDGFRβ adds further support to observations from the developmental biology literature that tendon fibroblasts are a distinct cell type from muscle fibroblasts (49).

*PDGFR inhibition prevents growth of tendon tissue after mechanical overload.* We next determined if inhibition of PDGFR signaling would impact the growth of tendons subjected to mechanical overload. To accomplish this, we used the small molecule CP-673,451 to block phosphorylation of both PDGFR subtypes in cultured tendon fibroblasts, and in whole tendon tissue. CP-673,451 inhibits both PDGFRα and PDGFRβ kinases with greater than 450-fold selectivity compared to other structurally related receptor tyrosine kinases, including vascular endothelial growth factor receptor (VEGFR), fibroblast growth factor receptor (FGFR), epidermal growth factor receptor (EGFR) and insulin-like growth factor 1 receptor (IGF1R) (43). Treatment of tendon fibroblasts with PDGF-BB resulted in phosphorylation of PDGFRα and PDGFRβ, and the addition of CP-673,451 blocked PDGFR phosphorylation (Figure 2A). In whole tendon tissue, mice treated with CP-673,451 demonstrated reduced levels of p-PDGFRα^Y849^/PDGFRβ^Y857^ after mechanical overload compared to vehicle-treated controls (Figure 2B). We also evaluated if kinases known to be downstream of PDGFR activation were impacted by CP-673,451 (38, 50), and observed significant attenuation of p-Akt^T308^ and p-ERK1/2^T202/Y204^ compared to controls (Figure 2B). These findings were also supported with histology, which also demonstrated CP-673,451 reduced PDGFR phosphorylation *in vivo* (Figure 2C).

**Figure 2.**
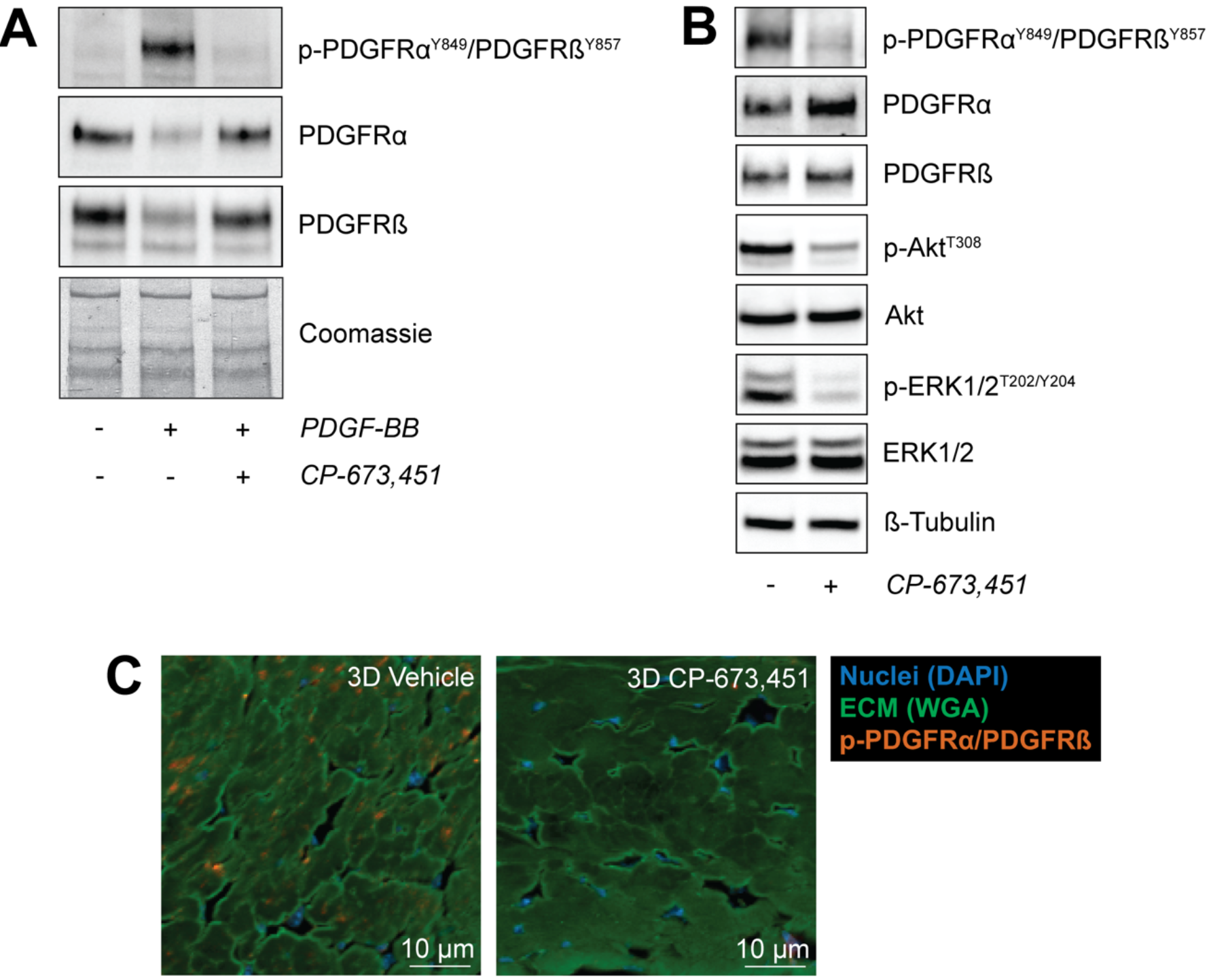
CP-673,451 inhibits the phosphorylation of PDGFRα and PDGFRβ in tendon fibroblasts in vitro and in vivo. (A) Representative immunoblots of serum-starved tendon fibroblasts treated with 20 ng/ml of PDGF-BB for 30 minutes with or without 1 μM of the PDGFR inhibitor CP-673,451. A Coomassie stained membrane is shown as a loading control. (B) Representative immunoblots of 3-day overloaded plantaris tendons demonstrating the ability of CP-673,451, to inhibit phosphorylation of PDGFRα and PDGFRβ in vivo. Phosphorylation of Akt and ERK1/2 were also inhibited by PDGFR inhibitor treatment. β-tubulin is shown as a loading control. (C) Immunohistochemistry of 3-day overloaded plantaris tendons treated with vehicle or PDGFR inhibitor showing a decrease in the abundance of p-PDGFRα/PDGFRβ-expressing cells in the overloaded plantaris tendons treated with CP-673,451 relative to vehicle-treated controls. Scale bars are 10 μm. DAPI, blue; WGA, green; p-PDGFRα^Y849^/PDGFRβ^Y857^, red.

We then explored morphological effects of PDGFR signaling on tendon growth. Mechanical overload of plantaris tendons resulted in outward growth of a neotendon matrix from the most superficial layers of the original tendon (Figure 3A), consistent with previous studies (14, 48). Overall, PDGFR inhibition did not impact the general morphological features of the plantaris tendons, but noticeable differences in tendon CSA and cell density were observed (Figure 3B-G). For the original tendon, the CSA of the 10-day vehicle group was slightly larger than both the 3-day vehicle and PDGFR inhibitor groups, but otherwise the CSA of the original tendon did not change in response to time after overload or treatment with PDGFR inhibitor (Figure 3B). In contrast, the CSA of the neotendon in the 3- and 10-day PDGFR inhibitor groups was 50% and 59% smaller compared to their respective vehicle-treated controls (Figure 3C). Changes in total tendon CSA generally followed the same trends observed in the neotendon data (Figure 3D). For the original tendon, the cell density of all groups was similar and did not change in response to time after overload or treatment with PDGFR inhibitor (Figure 3E). However, the cell density of the neotendon in the 10-day vehicle group was 60% higher compared to both the 3-day vehicle and PDGFR inhibitor groups, and this increase was not observed in response to PDGFR inhibition (Figure 3F). Similar to the total tendon CSA results, changes in total tendon cell density generally followed the same trends observed in the neotendon data (Figure 3G).

**Figure 3.**
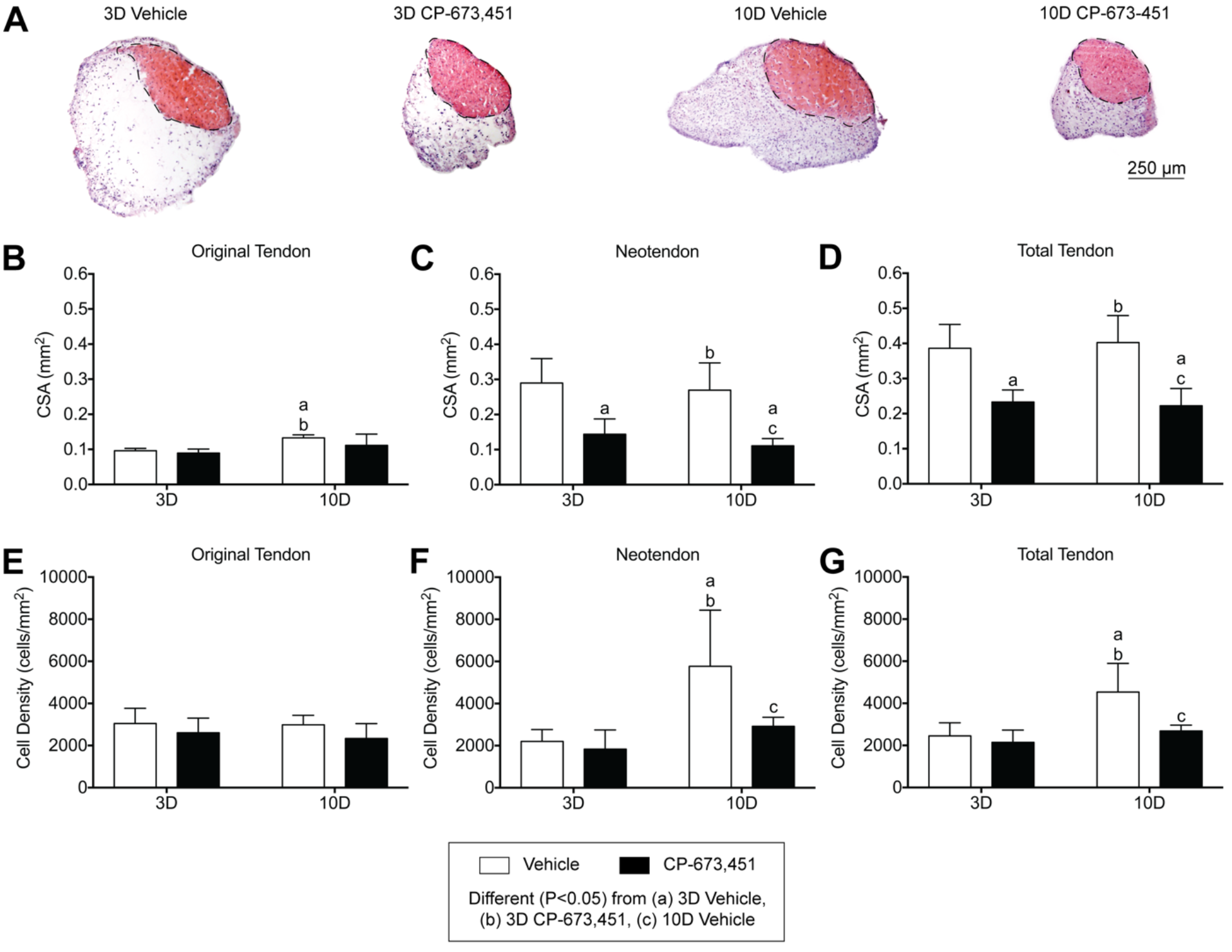
Inhibition of PDGFR signaling prevents growth of plantaris tendons subjected to mechanical overload. (A) Representative cross sections of 3- and 10-day overloaded plantaris tendons, treated with vehicle or PDGFR inhibitor, and stained with hematoxylin and eosin. Dashed black line indicates the boundary between the original tendon and neotendon. Scale bar is 250 μm. Quantitative analysis of (B-D) cross-sectional area (CSA, in mm^2^) and (E-G) cell density (cells/mm^2^) for the original tendon, neotendon and total tendon. Values are mean+SD. N=5 tendons for each group. Differences between groups were tested using a two-way ANOVA (α=0.05) followed by Tukey’s post hoc sorting: different (P<0.05) from a, 3D vehicle; b, 3D PDGFR inhibitor; c, 10D vehicle.

For cell proliferation, very few cells expressing the proliferation marker Ki67 were observed in the original tendon of mechanically overload tissue, and no difference between vehicle or PDGFR inhibitor groups were observed (Figure 4A-B). This is generally consistent with previous reports of the synergist ablation model in tendon, and supports the notion that tendon fibroblasts are largely a terminally differentiated cell population in adult tendons (14, 48). In contrast, the number of proliferating cells in the neotendon of the 3- and 10-day PDGFR inhibitor groups was 55% and 76% less than their respective vehicle-treated controls (Figure 4C). Changes in the number of proliferating cells in the total tendon generally followed the same trends observed in the neotendon (Figure 4D). Treatment of cultured tendon fibroblasts with PDGF-BB also increased cell proliferation by 33% (Figure 5A-B). Taken together, these results indicate that PDGFR inhibition suppresses the normal growth of tendon tissue after mechanical overload, and this is partly explained by defects in cell proliferation. The generally positive effects observed from the therapeutic use of PDGF in animal models of tendon injury may in part be due to the positive impact of PDGFR activation on tendon fibroblast proliferation (7, 29, 56).

**Figure 4.**
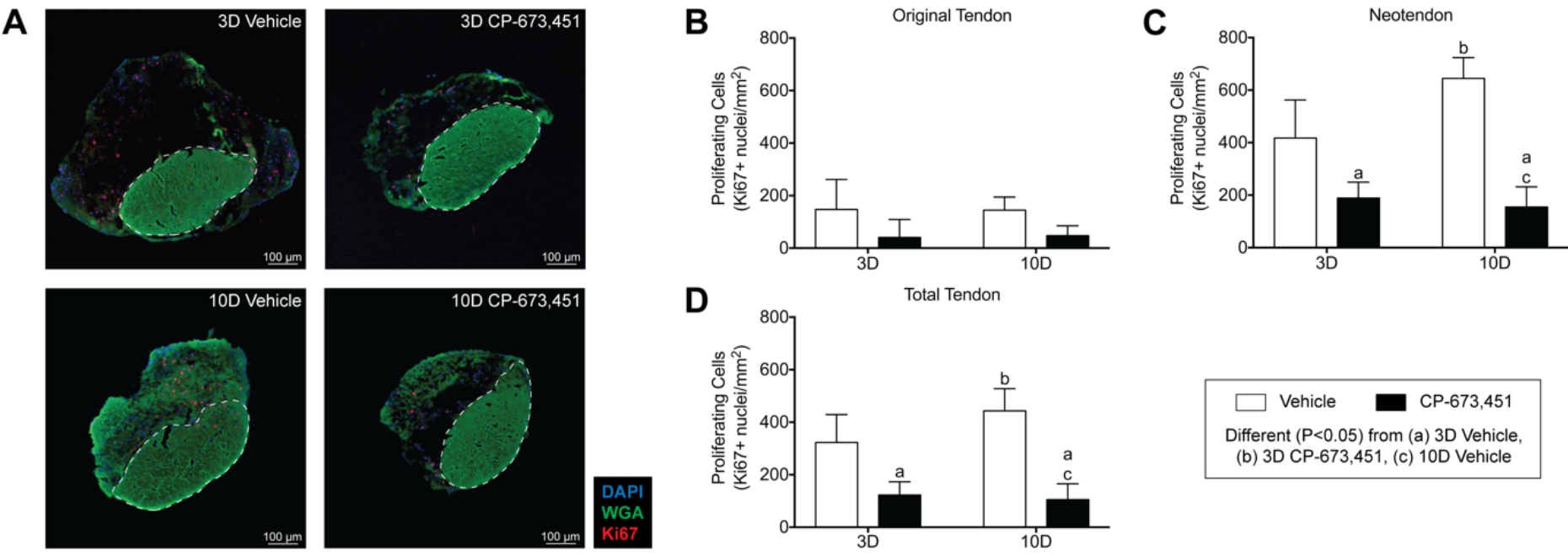
Inhibition of PDGFR signaling decreases tendon fibroblast proliferation in vivo. (A) Representative Ki67 immunostaining of 3- and 10-day overloaded plantaris tendons, treated with vehicle or PDGFR inhibitor. Dashed white line indicates the boundary between the original tendon and neotendon. Scale bars are 100 μm. DAPI, blue; WGA, green; Ki67, red. (B-D) Quantitative analysis of proliferating cells (Ki67^+^ nuclei/mm^2^) for the original tendon, neotendon and total tendon. Values are mean+SD. N≥4 tendons for each group. Differences between groups were tested using a two-way ANOVA (α=0.05) followed by Tukey’s post hoc sorting: different (P<0.05) from a, 3D vehicle; b, 3D PDGFR inhibitor; c, 10D vehicle.

*PDGFR-inhibited tendons demonstrate deficits in angiogenesis, ECM synthesis and remodeling, and cell specification and proliferation.* Since inhibition of PDGFR signaling resulted in reduced tendon growth, we next sought to further explore the mechanisms behind this response. We initially used microarray experiments and gene ontology (GO) analysis (Supplemental Figure 1) to identify genes to measure with qPCR. For markers of angiogenesis (Figure 6A), multiple genes demonstrated reduced expression in the 10-day PDGFR inhibitor compared to vehicle-treated controls, including platelet/endothelial cell adhesion molecule 1 (PECAM1), also known as CD31, which is a membrane glycoprotein involved in cell adhesion of endothelial cells, and CD146, another adhesion molecule that is highly expressed by pericytes (12). CD248, which is a C-type lectin domain protein important for angiogenesis (37), was reduced at 3 and 10 days in response to PDGFR inhibition. Thrombospondin 1 (TSP1), which is a secreted glycoprotein that inhibits angiogenesis through direct effects on endothelial cell proliferation and migration (18), was also downregulated at 10 days in the PDGFR inhibitor group compared to vehicle-treated controls, while slight changes in the expression of the angiogenic growth factor vascular endothelial growth factor A (VEGFA) were noted as a result of time after overload. These results are in general agreement with previous observations in skeletal muscle, where inhibition of PDGFR signaling prevented angiogenesis in mechanically overloaded muscles (50). Activation of PDGFRβ in pericytes appears important in the process of angiogenesis (42), and while we blocked both PDGFRs in this study and were not able to target PDGFRβ specifically, the reduction in angiogenesis-associated genes may be due to diminished pericyte activity.

**Figure 5.**
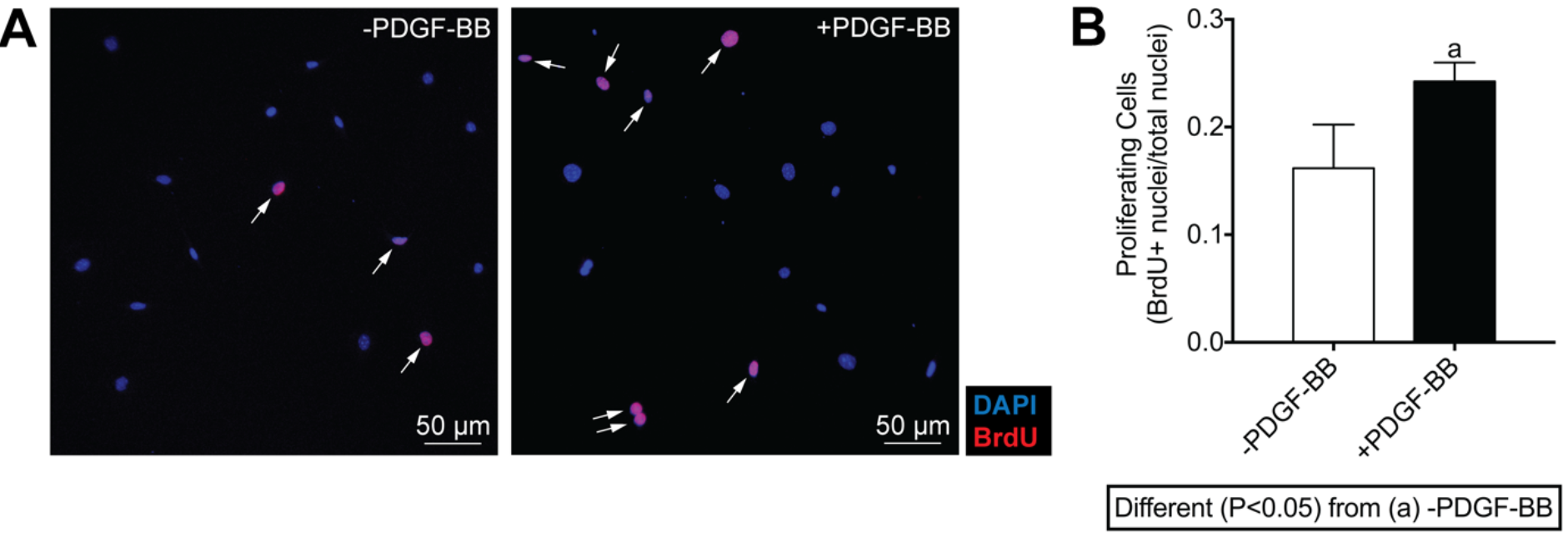
PDGF-BB stimulates proliferation of tendon fibroblasts in vitro. (A) *In vitro* proliferative activity of tendon fibroblasts was measured by BrdU uptake in the presence of 0.5% FBS with or without 20 ng/ml of PDGF-BB treatment. Proliferating cells were double labeled for DAPI (blue) and BrdU (red), and marked by white arrows. (B) Proliferative activity was quantified as mean±SD in 5 randomly selected fields of a single experiment from 6 independent experiments performed. Differences between groups were tested using an unpaired t-test; a, P<0.05.

**Figure 6.**
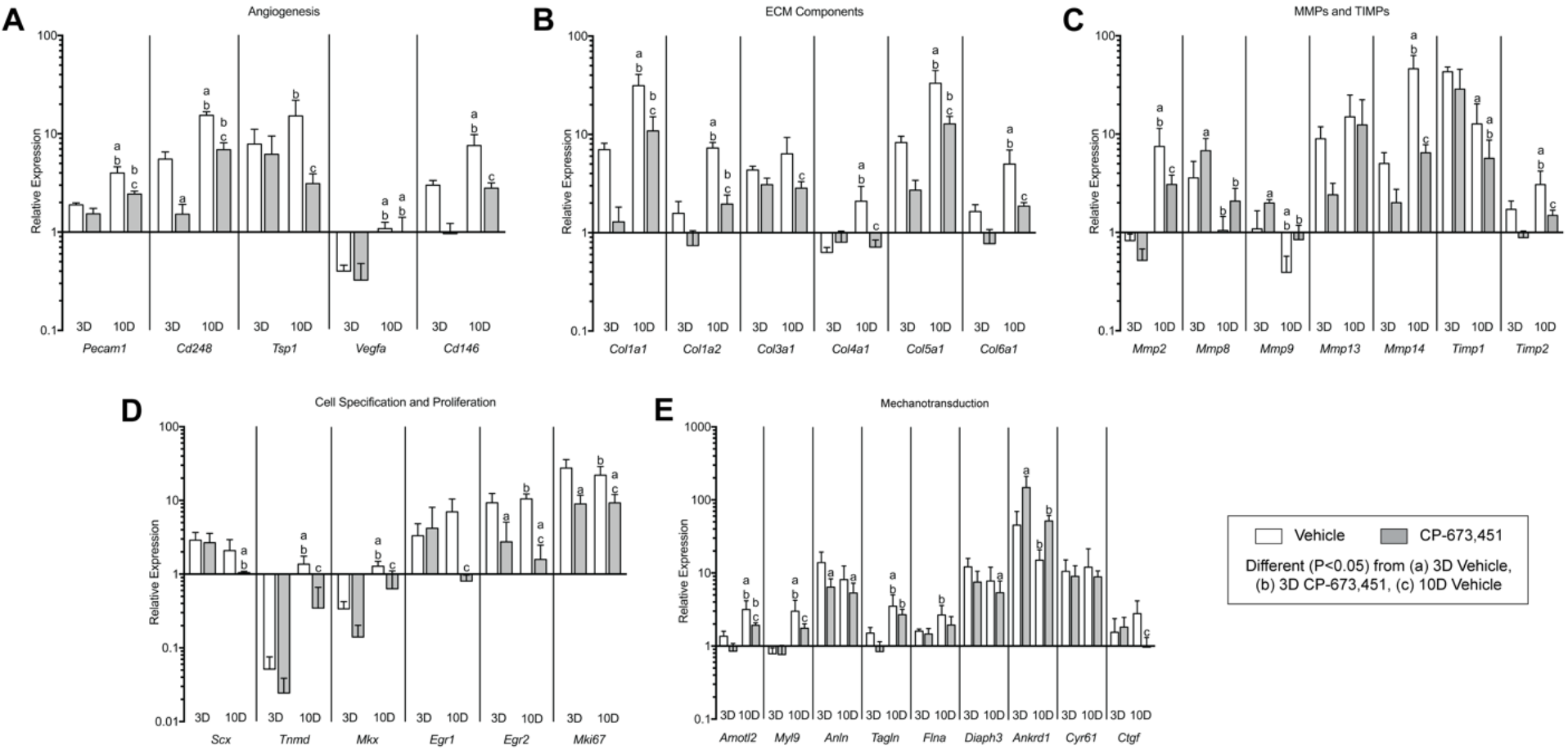
Inhibition of PDGFR signaling prevents the expression of angiogenesis, ECM synthesis and remodeling, cell specification and proliferation, and cell migration and mechanotransduction genes in plantaris tendons subjected to mechanical overload. Quantitative expression of (A) angiogenesis, (B) ECM synthesis, (C) MMPs and TIMPs, (D) cell specification and proliferation, and (E) mechanotransduction genes in 3- and 10-day overloaded plantaris tendons, treated with vehicle or PDGFR inhibitor. Target gene expression was normalized to the stable housekeeping gene peptidylprolyl isomerase D (PPID), and further normalized to plantaris tendons that were not subjected to synergist ablation. Values are mean+SD. N≥5 tendons for each group. Differences between groups were tested using a two-way ANOVA (α=0.05) followed by Tukey’s post hoc sorting: different (P<0.05) from a, 3D vehicle; b, 3D PDGFR inhibitor; c, 10D vehicle.

In addition to the changes observed in markers of angiogenesis, numerous ECM synthesis and remodeling genes were also affected by PDGFR inhibition (Figure 6B). Expression of the fibrillar types I, III and V collagens as well as the network types IV and VI collagens was not different at 3 days, but was reduced at 10 days in response to PDGFR inhibition. Given that type I collagen is the main structural protein of tendon, we sought to determine if the reduction in type I collagen transcripts at the whole tissue level was a direct effect of PDGFR inhibition on tendon fibroblasts, or an indirect effect related to a decrease in cell number. When cultured tendon fibroblasts were treated with PDGF-BB over a 12 hour time course, PDGF-BB stimulation had no significant effect on Col1a1 and Col1a2 transcript levels or pro-collagen 1a1 protein abundance (Figure 7A-C). This is consistent with findings from mechanically overloaded skeletal muscles, where reduced fibrillar collagen expression in PDGFR-inhibited animals occurred along with a reduction in markers of muscle fibroblast abundance (50). Combined, these results indicate that PDGF-BB does not regulate the expression of type I collagen at the mRNA and protein levels in tendon fibroblasts, and hence the reduction in type I collagen transcripts at the whole tissue level is likely an indirect effect on other signaling molecules that regulate type I collagen expression, or related to a decrease in cell number.

**Figure 7.**
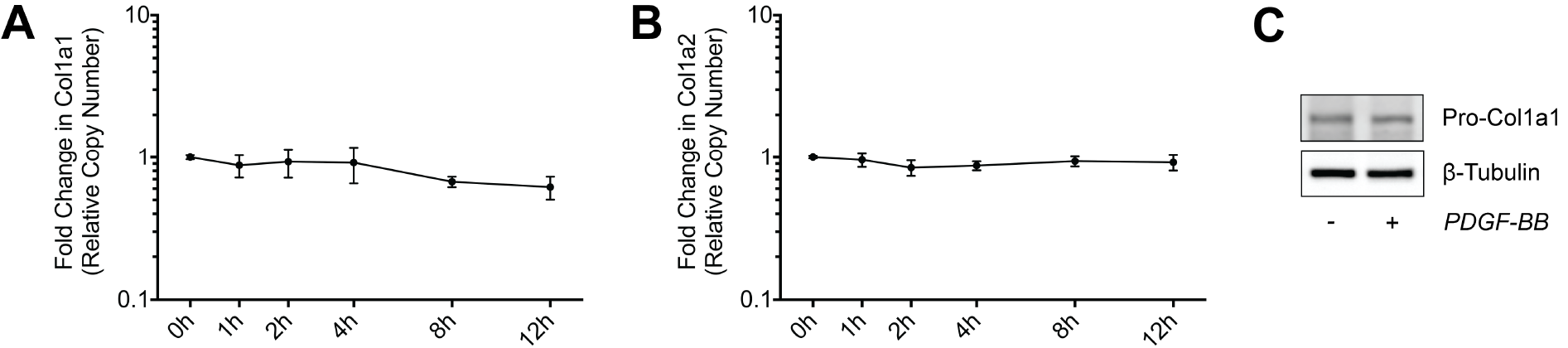
Effect of PDGF-BB treatment on collagen expression. Quantitative expression of (A) Col1a1 and (B) Col1a2 transcript levels in tendon fibroblasts treated with 20 ng/ml of PDGF-BB for 1, 2, 4, 8 and 12 hours. Values are mean±SD for N≥3 replicates. Differences between groups were tested using one-way ANOVA (P<0.05). No significant differences were found between groups. (C) Representative immunoblots of Pro-Col1a1 protein levels in tendon fibroblasts treated with 20 ng/ml of PDGF-BB for 24 hours in low-serum conditions. β-tubulin is shown as a loading control. Differences between groups were tested using a one-way ANOVA (α=0.05) followed by Tukey’s post hoc sorting.

Accompanying the changes observed in ECM synthesis and remodeling genes, PDGFR inhibition also resulted in marked changes in the expression of MMPs and TIMPs (Figure 6C). For the gelatinases, MMP2 expression was 59% lower at 10 days in the PDGFR inhibitor group compared to vehicle-treated controls, while MMP9 was 83% higher at 3 days in response to PDGFR inhibition. Similar to MMP9, the collagenase MMP8 was 89% higher at 3 days in response to PDGFR inhibition, whereas no differences in the expression of collagenase MMP13 were demonstrated in response to time after overload or treatment with PDGFR inhibitor. MT1-MMP, also known as MMP14, is a membrane tethered MMP that demonstrated the most profound upregulation in response to mechanical overload, and was also most substantially impacted by PDGFR inhibition. MT1-MMP was upregulated by greater than 5- and 46-fold at 3 and 10 days after mechanical overload, respectively, and treatment with PDGFR inhibitor prevented the increase in transcript levels at 10 days compared to vehicle-treated controls. Compared to the 3-day time point, TIMP1 transcript levels were lower in both 10-day groups, while TIMP2 transcript levels did not increase at 10 days in response to PDGFR inhibition. These findings are consistent with other studies that demonstrated PDGFR signaling was associated with increased expression and activation of MMPs, in particular MT1-MMP, as well as TIMPs, in other cell types (28, 45, 50, 52).

Transcription factors that play crucial roles in different stages of tendon development and specification (21) were upregulated after mechanical overload (Figure 6D). The basic helix-loop-helix transcription factor scleraxis (Scx) was upregulated greater than 2-fold at both 3 and 10 days and was not affected by PDGFR inhibitor treatment. After an initial downregulation of the atypical homeodomain transcription factor mohawk (Mkx) and the type II transmembrane glycoprotein tenomodulin (Tnmd) at 3 days, their transcript levels increased by 10 days compared to non-overloaded controls. The zinc finger transcription factors early growth response 1 and 2 (Egr1 and Egr2) did not increase at 10 days in response to PDGFR inhibition, while Egr2 expression was also 71% lower at 3 days in the PDGFR inhibitor group compared to vehicle-treated controls. Ki67 (Mki67) was reduced at 3 and 10 days in response to PDGFR inhibition. While pericytes are able to differentiate into cells which form blood vessels in a process dependent upon PDGFRβ activation, they are also able to differentiate into other cell lineages, including fibroblasts (12). Lineage tracing studies have not yet been performed, but there is compelling indirect evidence that pericytes are the progenitor cells of fibroblasts in adult tendon tissue (30, 48, 53), and it is possible that inhibiting PDGFR signaling resulted in reduced activity of fibroblast progenitor cells, which then decreased the expression of genes involved with tendon specification, proliferation, and differentiation.

There has been an association between the PDGF pathway and activation of the YAP signal transduction network in hepatic stellate cells (31), and we sought to evaluate whether PDGFR inhibition would impact the expression of YAP-regulated genes (Figure 6E) involved in mechanotransduction (4). Angiomotin-like protein 2 (AmotL2), which is a membrane-associated scaffold protein that localizes to lamellipodia during cell migration, and myosin light chain 9 (Myl9), which regulates contractility and cytoskeletal tension during cell migration, were downregulated at 10 days in response to PDGFR inhibition. The kinetic scaffold protein anillin (Anln), which binds F-actin and can recruit Rho GTPases to the leading edge of the cell membrane during cell migration, was reduced at 3 days in response to PDGFR inhibition. Other actin-binding proteins that help reorganize the actin cytoskeleton during cell migration include transgelin (Tagln), filamin A (Flna) and diaphanous related formin 3 (Diaph3), were not different between vehicle and inhibitor treated groups at each time point. In contrast, ankyrin repeat domain 1 (Ankrd1), which is a transcriptional repressor of MMP13, increased by greater than 3-fold in the 3-day PDGFR inhibitor group compared to vehicle-treated controls. The matricellular proteins cysteine-rich angiogenic inducer 61 (Cyr61) and connective tissue growth factor (CTGF) are key mediators of cell migration through their interaction with cell surface integrins, and while Cyr61 was not affected by PDGFR inhibition, CTGF expression was 65% lower at 10 days in the PDGFR inhibitor group compared to vehicle-treated controls. Overall, while several YAP-regulated genes were induced in overloaded tendons, only a few genes were differentially regulated between vehicle and PDGFR treated animals at a given time point. Although mechanotransduction mediated by YAP is likely important in controlling tendon growth, there does not appear to be marked interaction between the PDGFR and YAP pathways in mechanically overloaded tendons.

*MT1-MMP is an essential proteinase for fibroblast migration through reconstituted tendon ECM.* As MMPs were highly induced after mechanical overload and differentially regulated by PDGFR inhibitor treatment (Figure 6C), we sought to explore the role of PDGF-BB and MMPs in tendon fibroblast migration *in vitro*. Tendon fibroblasts were placed on top of a reconstituted collagen-rich gel reconstituted from tendon ECM, and exposed to growth factors and underwent manipulation of different kinases and effector proteins (Figure 8A). In the absence of specific growth factors, fibroblasts remained on the surface of the gel (Figure 8B-D). Treatment with transforming growth factor beta 1 (TGF-β1) did not cause fibroblasts to enter the gel, but did appear to increase proliferation as evidenced by a thicker layer of cells on the surface of the gel (Figure 8B-D). However, upon treatment with PDGF-BB, fibroblasts did migrate into the gel (Figure 8B-D). This process was prevented using the broad spectrum MMP inhibitor BB-94 (Figure 8B-D), indicating that migration was dependent upon MMP activity, which is in agreement with observations from other types of non-tendon cells (28, 45, 47, 52).

**Figure 8.**
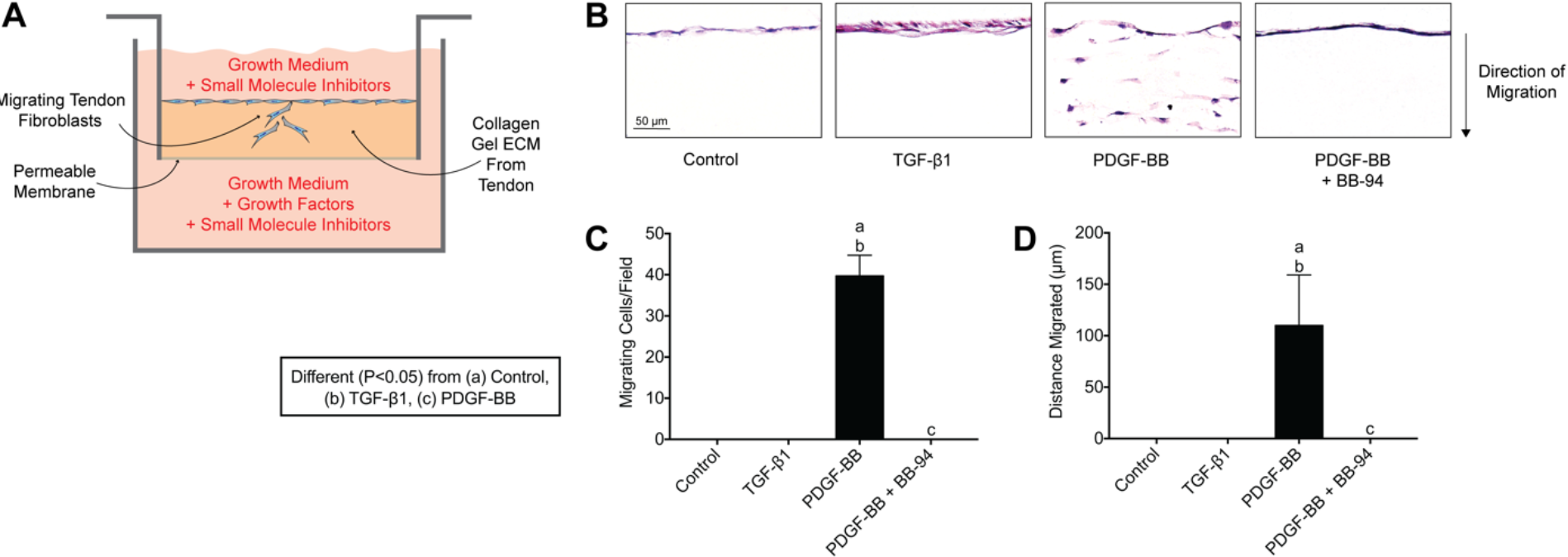
PDGF-BB stimulates tendon fibroblast migration in vitro. (A) Schematic representation of the *in vitro* migration assay. (B) Representative images of tendon fibroblasts obtained 6 days after being placed on the surface of a tendon ECM transwell system, treated with no additional growth factors (control), TGF-β1, PDGF-BB, and PDGF-BB + BB-94 (a synthetic broad spectrum MMP inhibitor). Quantification of (C) the number of migrating cells per field and (D) maximum distance migrated. Sections stained with hematoxylin and eosin. Values are mean+SD. N=4 replicates per group. Differences between groups were tested using a one-way ANOVA (α=0.05) followed by Tukey’s post hoc sorting: different (P<0.05) from a, control; b, TFG-β1; c, PDGF-BB.

As MT1-MMP was the most highly induced MMP after mechanical overload, and the expression of MT1-MMP was markedly attenuated in response to PDGFR inhibition at 10 days (Figure 6C), we then focused on the specific role of MT1-MMP in tendon fibroblast migration. Knocking down MT1-MMP expression with siRNA prevented the migration of fibroblasts into the tendon gel (Figure 9A-C). Taken together, these results indicate that MT1-MMP is required for the migration of tendon fibroblasts through their ECM.

**Figure 9.**
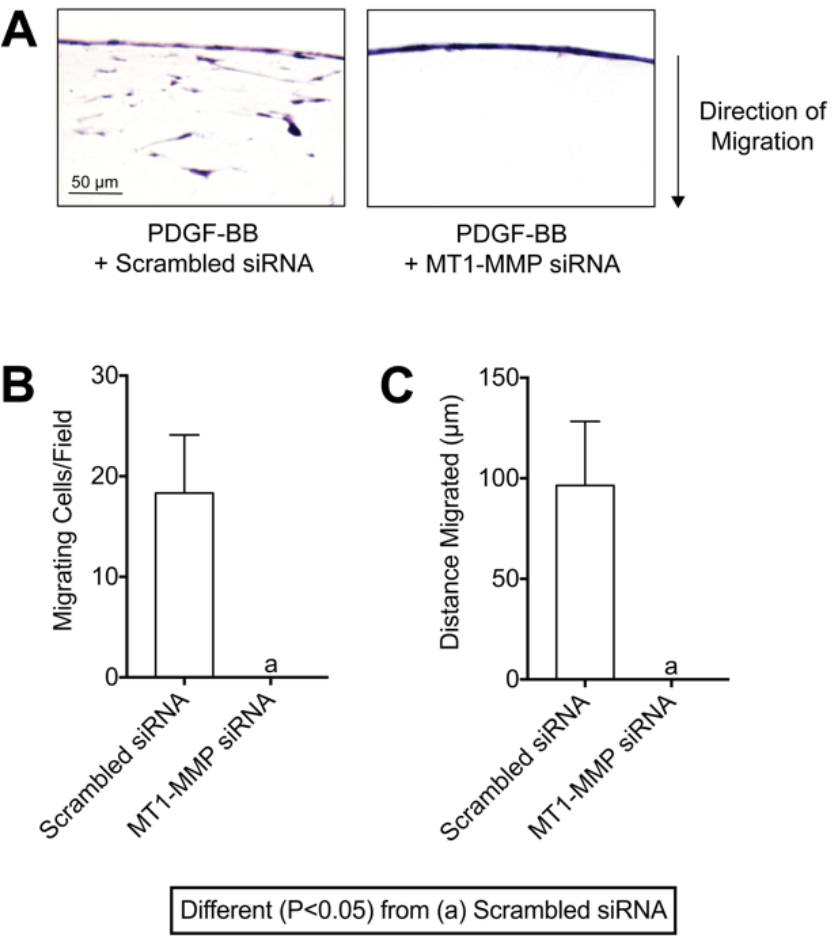
MT1-MMP is required for PDGF-BB-mediated fibroblast migration through tendon ECM. (A) Representative images of tendon fibroblasts transfected with scrambled siRNA or MT1-MMP siRNA, and treated with PDGF-BB, measured after 6 days. Quantification of (B) the number of migrating cells per field and (C) maximum distance migrated of the MT1-MMP-silenced tendon fibroblasts. Sections stained with hematoxylin and eosin. Values are mean+SD. N=4 replicates per group. Differences between groups were tested using an unpaired Student’s t-test; a, P<0.05.

*PDGF-BB controls the translation, membrane targeting and activation of MT1-MMP in tendon fibroblasts.* As PDGF signaling directed tendon fibroblast migration in a MT1-MMP dependent fashion, we next sought to identify the signaling pathways which regulate this process by evaluating the phosphorylation of various kinases that have previously reported to be involved in migration and proliferation of other cell types (54). In response to activation of the PDGFRs, there was a rapid phosphorylation of ERK1/2 and Akt, followed soon thereafter by p70S6K which is downstream from Akt (Figure 10A). We then stimulated tendon fibroblasts in transwell systems with PDGF-BB and either PD98059 to block ERK1/2 phosphorylation through inhibition of its upstream kinase MEK1 (40), or Wortmannin to block Akt phosphorylation through inhibition of its upstream kinase PI3K (2). In response to treatment with either of these inhibitors, there was a greater than 90% reduction in the number of migrating cells per field, and more than 80% in the distance migrated by these cells (Figure 10B-C).

**Figure 10.**
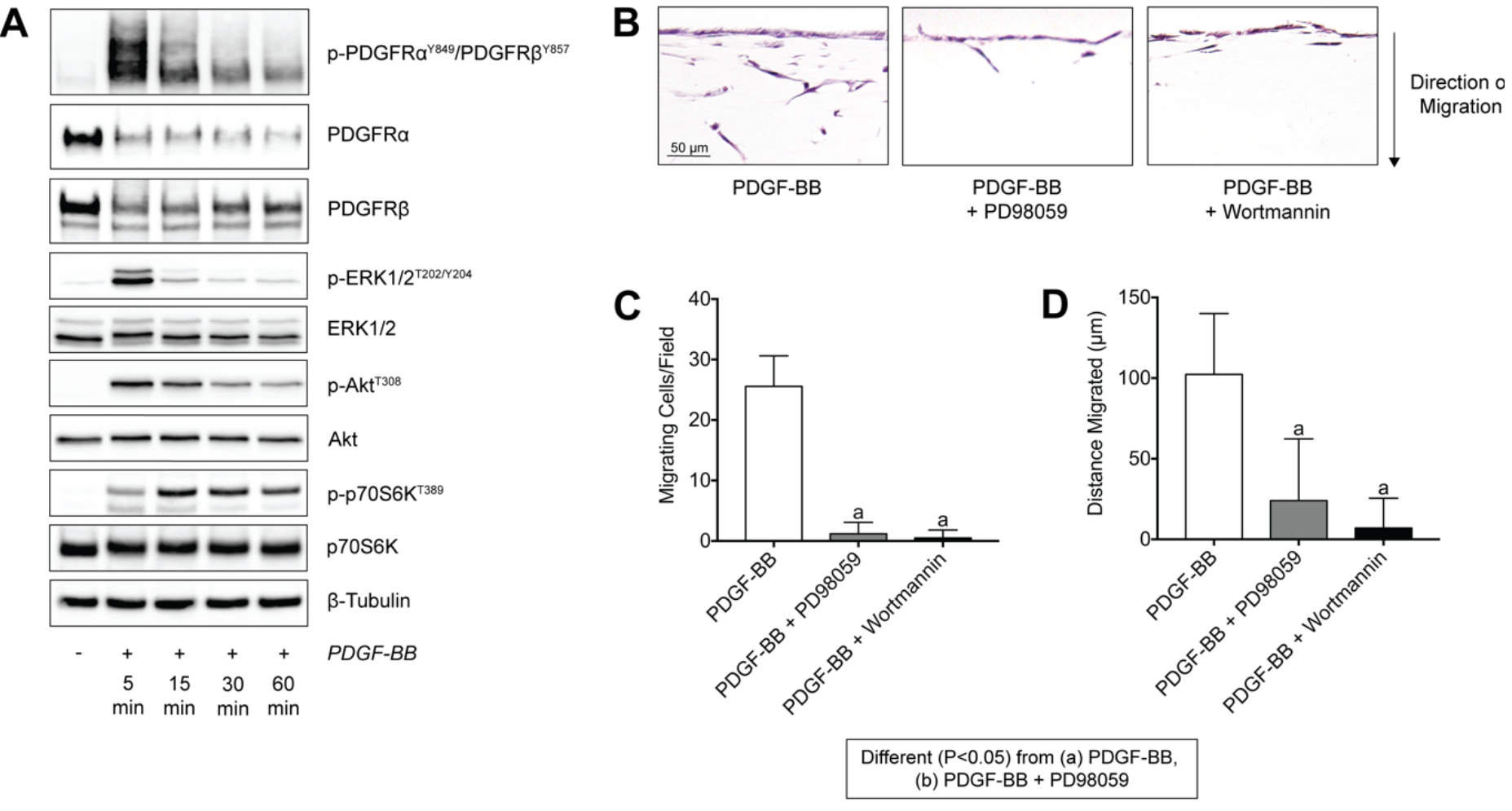
PDGF-BB stimulation of tendon fibroblasts activates PI3K/Akt and ERK1/2 pathways, which both in turn mediate tendon fibroblast migration through tendon ECM. (A) Representative immunoblots of total and phospho PDGFRα, PDGFRβ, ERK1/2, Akt and p70S6K from serum-starved tendon fibroblasts treated with PDGF-BB for 5, 15, 30 and 60 minutes. β-tubulin is shown as a loading control. (B) Representative images of tendon fibroblasts obtained 6 days after being placed on the surface of a tendon ECM transwell system, treated with PDGF-BB, PDGF-BB + PD98059, or PDGF-BB + Wortmannin. Quantification of (C) the number of migrating cells per field and (D) maximum distance migrated. Sections stained with hematoxylin and eosin. Values are mean+SD. N=4 replicates per group. Differences between groups were tested using a one-way ANOVA (α=0.05) followed by Tukey’s post hoc sorting: different (P<0.05) from PDGF-BB; b, PDGF-BB + PD98059.

PDGF-BB did not change the transcript levels of MT1-MMP in cultured tendon fibroblasts (Figure 11A), and we therefore evaluated how PDGF-BB signaling regulated MT1-MMP in a post-transcriptional fashion. The specificity and efficacy of PD98059 and Wortmanin to block ERK1/2 and Akt signaling, respectively, was verified (Figure 11B-D). MT1-MMP is produced in an inactive 63kD pro-form, that is eventually activated as a 60kD form, and after some time is cleaved to a 45kD inactive, processed form (8). The treatment of tendon fibroblasts with PDGF-BB increased total MT1-MMP protein levels, including pro-MT1-MMP, active MT1-MMP and processed MT1-MMP (Figure 11E-H). Inhibiting ERK1/2 resulted in a further increase in pro-MT1-MMP, but reduced active and processed MT1-MMP (Figure 11E-H). Blocking Akt reduced total MT1-MMP levels, which is consistent of the role of the downstream kinase p70S6K in phosphorylating the ribosomal protein S6 that initiates protein synthesis by ribosomes (13). Additionally, inhibiting Akt did not change pro-MT1-MMP levels compared to PDGF-BB stimulation alone, but did decrease active and processed MT1-MMP (Figure 11E-H). When the ratio of active to pro-MT1-MMP was calculated, inhibition of ERK1/2 resulted in a reduction in active MT1-MMP relative to pro-MT1-MMP compared to other treatments (Figure 11I).

**Figure 11.**
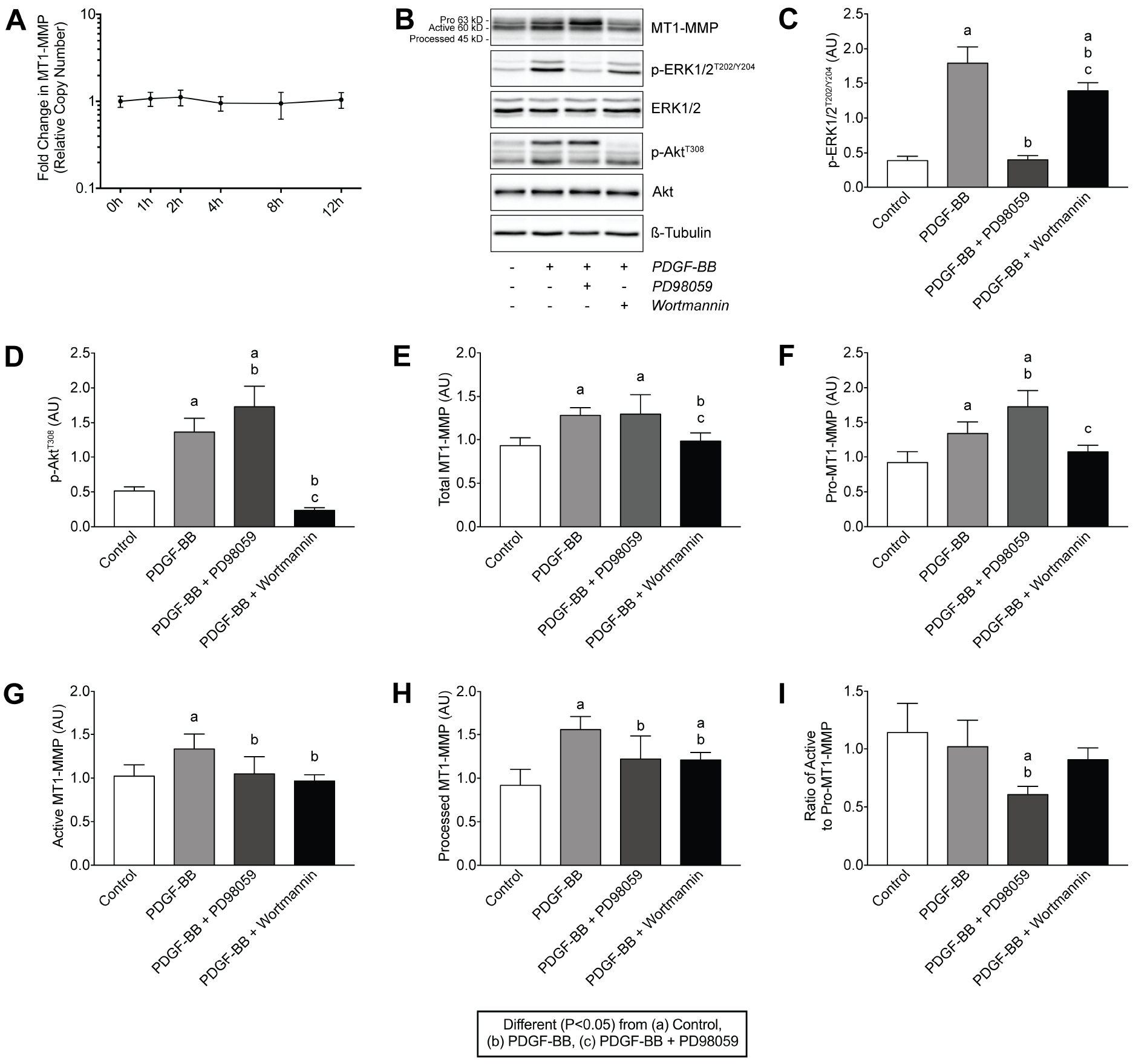
PDGF-BB does not regulate MT1-MMP mRNA expression, but does increase MT1-MMP protein levels through a PI3/Akt-dependent mechanism. (A) Quantification of MT1-MMP transcript levels in tendon fibroblasts treated with 20 ng/ml of PDGF-BB for 1, 2, 4, 8 and 12 hours. Values are mean±SD for N≥3 replicates. Differences between groups were tested using one-way ANOVA (P<0.05). (B) Representative immunoblots of Pro-MT1-MMP (63 kD), active MT1-MMP (60 kD), processed MT1-MMP (45 kD), and phospho and total ERK1/2 and Akt are shown, from tendon fibroblasts incubated alone or with PDGF-BB in the presence or absence of PD98059 or Wortmannin for 24 hours. β-tubulin is shown as a loading control. Quantification of (C) p-ERK1/2^T202/Y204^, (D) p-Akt^T308^, (E) total MT1-MMP, (F) pro-MT1-MMP, (G) active MT1-MMP and (H) processed MT1-MMP protein levels. (I) Ratio of the pro- to active form of MT1-MMP. Values are mean±SD for 6 replicates. Differences between groups were tested using a one-way ANOVA (α=0.05) followed by Tukey’s post hoc sorting: a, different (P<0.05) from control; b, PDGF-BB; c, PDGF-BB + PD98059.

Finally, because MT1-MMP exerts its effects in the extracellular matrix, we determined whether ERK1/2 or Akt regulated the trafficking of MT1-MMP to the plasma membrane.
Inhibition of both kinases prevented the PDGF-BB-induced localization of MT1-MMP to the plasma membrane (Figure 12A-C). Taken together, these results indicate that in tendon fibroblasts, PDGF-BB controls the translation of MT1-MMP mRNA through the Akt/p70S6K pathway, ERK1/2 is important for the activation of MT1-MMP, and that the trafficking of MT1-MMP to the plasma membrane is dependent upon both ERK1/2 and Akt signaling pathways (Figure 12D). These findings differ from previous reports in bone marrow-derived mesenchymal stem cells which indicated that treatment with PDGF-BB increased MT1-MMP mRNA expression and protein levels in a PI3K/Akt- and ERK1/2-dependent manner (52), suggesting that the regulation of MT1-MMP expression in response to PDGF-BB stimulation is cell type and tissue specific.

**Figure 12.**
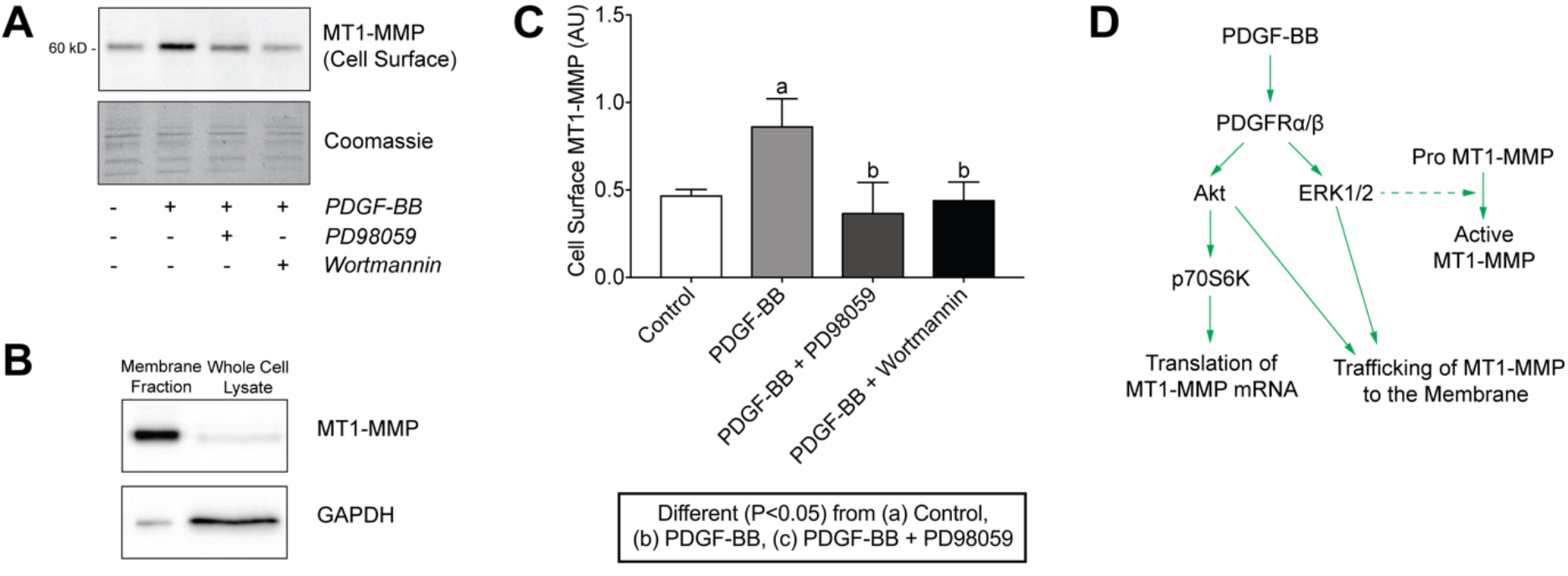
PDGF-BB-dependent membrane trafficking of MT1-MMP in tendon fibroblasts is regulated by the PI3K/Akt and ERK1/2 pathways. (A) Representative immunoblots of cell surface MT1-MMP protein levels in tendon fibroblasts after incubation alone or with PDGF-BB in the presence or absence of PD98059 or Wortmannin for 24 hours in low-serum conditions. Coomassie staining is shown as a total protein loading control. (B) Representative immunoblots of the membrane fraction and whole cell lysates of tendon fibroblasts using MT1-MMP and GAPDH as typical membrane and cytosolic proteins, respectively. Due to the isolation detergents, membrane fraction lanes run in a narrow fashion. (C) Quantification of cell surface MT1-MMP protein levels. Values are mean±SD for N≥3 replicates. Differences between groups were tested using a one-way ANOVA (α=0.05) followed by Tukey’s post hoc sorting: a, different (P<0.05) from control; b, PDGF-BB; c, PDGF-BB + PD98059. (D) Diagram of PDGF-BB-dependent regulation of MT1-MMP expression and membrane trafficking in tendon fibroblasts.

*Limitations.* There are several limitations to the current study. We used a specific inhibitor of PDGFRα and PDGFRβ (43), and were not able to selectively inhibit each receptor to determine its individual function. Although more than 20 MMPs have been described, we only targeted MT1-MMP during our migration assays, and it remains possible that other MMPs are also important for tendon fibroblast migration. Additionally, we measured differences in tendon morphology after mechanical overload due to PDGFR inhibition, but tendon mechanics and other functional assays were not performed. The synergist ablation procedure is reliable, reproducible and is well tolerated by mice, but the net mechanical load placed on the plantaris tendon is greater than that experienced during normal locomotion or as a result of treadmill training. Furthermore, while our study focused only on the plantaris tendon, the observed differences might not be reflected in other trunk or limb tendons. We measured the expression of many transcripts using microarray and qPCR, but changes in the transcriptome might not follow changes in the proteome. Finally, we collected data at only two time points after mechanical overload, and additional time points are likely to provide further mechanistic insight. Despite these limitations, we feel this work provides an important contribution to our understanding of how PDGFR signaling controls tendon growth in adult animals.

*Summary and Future Directions.* In response to changes in mechanical load, tendon actively remodels its ECM to meet the new demands placed on it, which is typically accompanied by increases in tendon CSA, cell density, and type I collagen content (14, 26, 34). Tendon fibroblasts constantly sense and respond to biomechanical and biochemical cues in their environment, and an increase in mechanical load placed on the tendon is often a signal for growth (15, 59). In the current study, we report that tendon fibroblasts express PDGFRα and PDGFRβ, that PDGFR signaling is required for the load-induced growth of tendons in adult animals, and MT1-MMP is an essential proteinase for the migration of tendon fibroblasts through their ECM via Akt and ERK1/2 dependent mechanisms. While PDGFR inhibition resulted in reduced MT1-MMP expression at the whole tissue level, treatment of tendon fibroblasts with PDGF-BB directly did not alter MT1-MMP expression, suggesting the results observed at the whole tissue level are likely due to indirect effects. These combined findings help to provide mechanistic insights into the generally positive outcomes in applied studies that have used PDGF-BB to stimulate tendon regeneration (7, 29, 56), and the disordered ECM observed in the tendons of mice which lack MT1-MMP in tendon fibroblasts (ScxCre:MT1-MMP^f/f^) (55). Increased PDGFRβ expression and greater proliferation has been observed in biopsies of patients with patellar tendinopathy compared to healthy controls (44), and it is possible that the targeted manipulation of PDGFR signaling may be useful in the treatment of chronic tendon disorders.

Apart from its role in cell migration, MT1-MMP also regulates other key cellular processes including proliferation, differentiation and cell survival, by either changing the tissue architecture or converting labile matrix proteins into active signaling molecules (23). Although the mechanisms underlying the ERK1/2-mediated activation of MT1-MMP are not known, furin is a protease that is a member of the subtilisin-like proprotein convertase (SPC) family and is known to activate MT1-MMP (23). Members of the SPC family are known to be regulated by PDGF-BB (3), and furin expression has been shown to be regulated by ERK1/2 activation (41). Additional studies of furin and other SPC proteases are likely to provide further insight into the molecular and biochemical mediators of tendon growth in adult animals.

## ACKNOWLEDGEMENTS

The authors would like to acknowledge helpful discussions and technical support by Dr. Stephen Weiss and Dr. Farideh Sabeh.

## GRANTS

This work was supported by NIH/NIAMS grants R01-AR063649 and F32-AR067086.

## DISCLOSURES

The authors declare they have no conflicts of interest with the contents of this article.

## AUTHOR CONTRIBUTIONS

KS and CM designed the study; KS, JM, ND, AR, DS performed the research; KS, JM, ND, DS and SB analyzed the data; and KS and CM wrote the paper; CM supervised the study.

